# Conspecific sperm precedence is reinforced but sexual selection weakened in sympatric populations of *Drosophila*

**DOI:** 10.1101/071886

**Authors:** Dean M. Castillo, Leonie C. Moyle

## Abstract

Sexual selection is well recognized as a driver of reproductive isolation between lineages. However, selection for increased reproductive isolation could reciprocally change the outcomes of sexual selection, when these processes share a genetic basis. Direct selection for reproductive isolation occurs in the context of ‘reinforcement’, where selection acts to increase prezygotic barriers to reduce the cost of heterospecific matings. Many studies of reinforcement focus on premating reproductive barriers, however postmating traits-such as conspecific sperm precedence (CSP)-can also respond to reinforcing selection. We tested whether i) CSP responded to reinforcing selection, and ii) this response in sympatric populations altered intraspecific sperm competition (ISC) and the strength of sexual selection, with the sister species *Drosophila pseudoobscura* and *D. persimilis*. We used sperm competition experiments to evaluate differences in CSP and ISC between two sympatric and two allopatric populations of *D. pseudoobscura*. Using multiple genotypes for each population allowed us to estimate not only patterns of phenotype divergence, but also the opportunity for sexual selection within each population. Consistent with a pattern of reinforcement, the sympatric populations had higher mean CSP. Moreover, ISC was altered in sympatric populations, where we observed decreased average offensive sperm competitive ability against conspecific males, allowing less opportunity for sexual selection to operate within these populations. These data demonstrate that strong reinforcing selection for reproductive isolation can have consequences for sexual selection and sexual interactions within species, in these important postmating sperm competition traits.

## Introduction

When closely-related species comes into contact, the presence of heterospecifics can influence sexual interactions and therefore alter patterns of selection on reproductive traits. In cases where these species have the potential to interbreed, selection can favor divergence in sexual traits to avoid costs of heterospecific mating, a type of reproductive character displacement commonly called reinforcement [1–3]. The frequency at which reinforcement contributes to speciation is still under debate [3–4] although several recent examples provide strong evidence for reinforcement acting on mating traits [5–10]. Regardless, the mating trait changes that evolve in response to reinforcement can have collateral effects on intraspecific sexual dynamics [6]. This can in turn alter the magnitude and efficacy of sexual selection specifically within populations exposed to heterospecifics. These potential reciprocal interactions between sexual selection and reproductive isolation remain relatively untested [6–7], but can have important consequences for how we interpret evolution of sexual traits and interactions. For example, patterns of reproductive trait evolution in rapid radiations, where sexual selection is thought to be the primary driver, may be misinterpreted if they do not take into account species interactions.

For reinforcement and sexual selection to reciprocally affect the evolution of sexual traits, these traits must be involved in both processes and share a genetic basis. Currently the best example of a shared genetic basis for sexual selection and reproductive isolation comes from *Drosophila* sperm competition genes, several of which have been shown to mediate both sexual selection through intraspecies sperm competition (ISC) and reproductive isolation via conspecific sperm precedence (CSP) [11]. Conspecific sperm precedence occurs when a female mates with both heterospecific and conspecific males yet most of the progeny are sired by the conspecific male; this precedence can occur either through competitive mechanisms (including male sperm competition and cryptic female choice) or non-competitive mechanisms (resulting mainly from gametic incompatibilities). CSP has proven to be a strong reproductive isolation barrier among species in *Drosophila* [4,12–13] and in many other plant and animal species [4, and references therein]. Although ubiquitous, CSP can be overlooked as a reproductive isolating barrier because it involves inconspicuous phenotypes that are not readily observed in the field [14]. Moreover, although reinforcement studies have overwhelmingly focused on pre-mating traits, postcopulatory prezygotic traits including CSP can also be the target of reinforcement [15–17]. Previous empirical studies have been equivocal about whether heterospecific interactions and reinforcement select for increased CSP specifically in sympatry, with no single study simultaneously estimating and comparing levels of CSP in allopatric and sympatric populations [13, 18–25]

While reinforcing selection (acting on CSP) and sexual selection (acting on ISC) could interact to influence evolutionary change in post-copulatory traits, the outcomes of this interaction clearly will depend upon whether these forces act in concert or in opposition. When sexual selection and reinforcing selection act in concert, trait evolution can proceed faster than otherwise expected, but the direction of trait evolution remains unchanged. In contrast, the potential feedback between sexual selection and reproductive isolation can generate complex evolutionary outcomes when these forces act at cross-purposes. For example, sperm competition is shaped by sexual conflict between males and females (i.e. antagonistic pleiotropy [26–28]) and genotype-genotype interactions (male-male [29–30] and male-female: [31–33]. Both are expected to maintain high variance in the affected traits and, indeed, sperm competition genes are often highly variable both in terms of molecular and phenotypic variation [30, 33–34]. In contrast, under models of speciation by sexual selection—where isolation is generated by strong disruptive selection between populations and directional selection within a population [35–36]—genetic variance of traits that act as barriers to reproduction is expected to be reduced and the overall trait mean shifted. The net effect of selection imposed by intrapopulation sexual interactions and by reinforcement can together produce phenotypic and genetic variation in sperm competition traits/genes that is different from the optimal variation when sexual selection acts alone.

One way these potentially antagonistic optima could play out is when reinforcement-mediated changes in the mean and variance of sperm competition traits alter the opportunity for sexual selection among conspecifics [7]. Sperm competition contributes to variance in reproductive success because male genotypes that can disproportionately sire offspring increase their fitness compared to the fitness of rival male conspecifics [37–38]. Strong sperm competition leads to greater opportunity for sexual selection because there is greater variance in reproductive success compared to scenarios where males have equal probability of siring offspring. This generates two alternative predictions of the possible effects of reinforcement on sexual selection. First, the response to strong directional selection from reinforcement on sperm competition traits could lead to greater siring ability in intrapopulation sperm competition, increasing variance in reproductive success and opportunity for sexual selection. Alternatively, strong directional selection could reduce phenotypic variation so that competitive ability is equalized among males, thus reducing the opportunity for sexual selection.

The strategy we used to evaluate the interaction between selection for increased reproductive isolation (i.e. reinforcement) and sexual selection acting on sperm competition genes was to estimate variation between genotypes in CSP and ISC in parallel. Both CSP and ISC are measures of postcopulatory offensive sperm competition, estimated by allowing females to mate sequentially with two different male genotypes and scoring the paternity of the resulting progeny. Here our focus was on second-male or ‘offensive’ siring success. This is typically referred to as ‘P2’ and captures the ability of the second mated male to sire offspring by displacing or disabling the sperm of the first male. For our experiments the first male was either heterospecific (to estimate CSP) or conspecific (to estimate ISC) tester male. By comparing the relative competitive success of replicate male lines against a common set of either heterospecific and conspecific male tester genotypes, we could estimate post-copulatory CSP and ISC in parallel in the same experiment. Using this design we also estimated which genotype effects (male genotype, female genotype, or the interaction) might shape CSP and ISC. Females experience the most cost of heterospecific matings [39–41] and could control CSP via cryptic female choice [42], thus we would expect strong female genotype effects on CSP. This contrasts with previous studies of ISC where both male and female genetic effects, and their interaction were significant effects [31,33]. Unlike ISC the phenotypic and genetic variance for CSP has not been empirically explored and their similarity to ISC is currently unknown.

In this study, we examine evidence for reinforcement of CSP among populations of *Drosophila pseudoobscura* that are allopatric or sympatric with their closely related sister species *D. persimilis*, and evaluate the potential consequences of these heterospecific interactions for ISC and sexual selection within *D. pseudoobscura* populations. One of the first clear empirical demonstrations of reinforcement on premating isolation was described in this species pair [43]. This finding suggests that heterospecific interactions and matings are frequent and sustained over evolutionary time and can act as a substantial selective agent on reproductive traits in this system. Here we determine whether there is evidence that heterospecific interactions have selected for increased CSP, by comparing this barrier among populations of *D. pseudoobscura* that are allopatric or sympatric with *D. persimilis*. A pattern of stronger CSP specifically in sympatry is consistent with reinforcement; moreover, because postcopulatory traits are less likely to be directly affected by environmental conditions, this pattern is unlikely to be explained by alternative phenomena, such as ecological selection, that could also explain character displacement in sympatry (see Discussion). Using a consistent design across all populations we could also estimate premating reproductive isolation in the same experiment, and compare its strength in sympatry and allopatry. Second, we evaluate whether selection for strong CSP in sympatry has affected ISC, and thereby post-copulatory sexual selection, as might occur when CSP and ISC have shared genetic architecture. Throughout, we test for differences in trait variation across a set of distinct genotypes which allows us to specifically evaluate which sex is playing a more critical role in determining variation in heterospecific and conspecific postcopulatory interactions.

## RESULTS

### No difference between allopatric and sympatric populations in premating isolation

Because our experimental assessment of CSP involved first mating with a heterospecific *D. persimilis* male, we were able to estimate the magnitude of premating isolation in each *D. pseudoobscura* population in our experiment. We did not find evidence for a pattern consistent with reinforcement of premating isolation mediated by female mate preference. The average probability of heterospecific matings ranged from 46-52% between populations, and did not differ between allopatric and sympatric populations (χ^2^ test of independence: χ^2^=1.185, df=1, *P*=0.2763; Wald’s Test: χ^2^=19, df=4, *P*=0.75; Table 1). In pairwise tests between each allopatric and sympatric population we also failed to reject the null hypothesis. Though we did not detect a signal of reinforcement there was ample genetic variance in heterospecific mating rate between female genotypes available for selection within each population ((Fig. 1; Supplemental Table 1). Only in one of the populations (Lamoille, which is allopatric) did the identity of the *D. persimilis* tester line affect variation in premating isolation (Supplemental Table 1).

**Table 1.** The average levels of reproductive isolation for each *D. pseudoobscura* population measured from two barriers to reproduction: female preference (proportion of females that did not mate with heterospecifics) and conspecific sperm precedence (CSP). Higher values indicate stronger reproductive isolation. Interpopulation sperm precedence (ISC) is included for comparison. The mean and variance estimates for CSP and ISC are based on 64 replicates per populations A = allopatric; S = sympatric

|  | Fem. Pref. | CSP |  | ISC |  |
| --- | --- | --- | --- | --- | --- |
| Population | Proportion (n) | Mean | Variance | Mean | Variance |
| Lamoille (A) | 0.481 (179) | 0.75 | 0.054 | 0.76 | 0.028 |
| Zion (A) | 0.476 (145) | 0.77 | 0.041 | 0.80 | 0.047 |
| Mt. St. Helena (S) | 0.540 (200) | 0.90 | 0.017 | 0.79 | 0.057 |
| Sierra (S) | 0.505 (222) | 0.92 | 0.018 | 0.68 | 0.052 |

**Figure 1.**
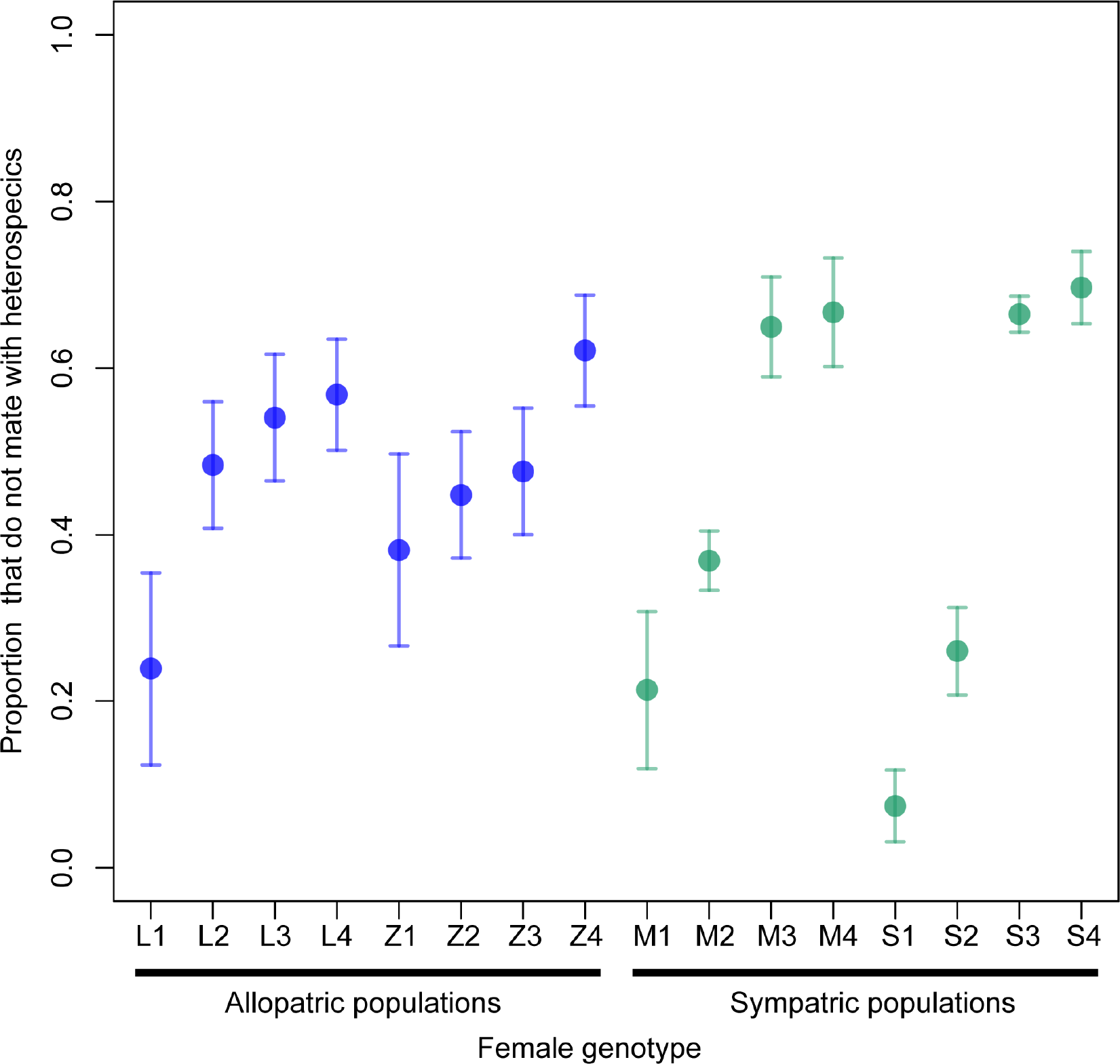
Prezygotic reproductive isolation via female mating preference does not show a pattern consistent with reinforcement. Reproductive isolation is measured by the proportion of females that did not mate with heterospecifics in individual no-choice trials. Significant variation among *D. pseudoobscura* female genotypes in female preference occurs in each population (Supplemental Table 1). Each point is the mean reproductive isolation for each isofemale line tested against each of four *D. persimilis* tester males. Error bars represent ± one standard error.

### Reinforcement acts on conspecific sperm precedence

Unlike premating isolation, we observed a pattern consistent with reinforcement for conspecific sperm precedence (CSP). Specifically, in sympatry we find both greater average CSP (*t*=−6.5898, df=210.92, *P*<0.001; Wilcox *W*=4427.5, *P*<0.001) and less phenotypic variation in this trait (Levene-type test χ^2^=22.82, *P*<0.0001) when data were pooled by geographic region (allopatry versus sympatry) (Table 1; Fig. 2A). These differences in both the average and variance of CSP were also observed in pairwise tests between individual allopatric and sympatric populations (Supplemental Table 2).

**Figure 2.**
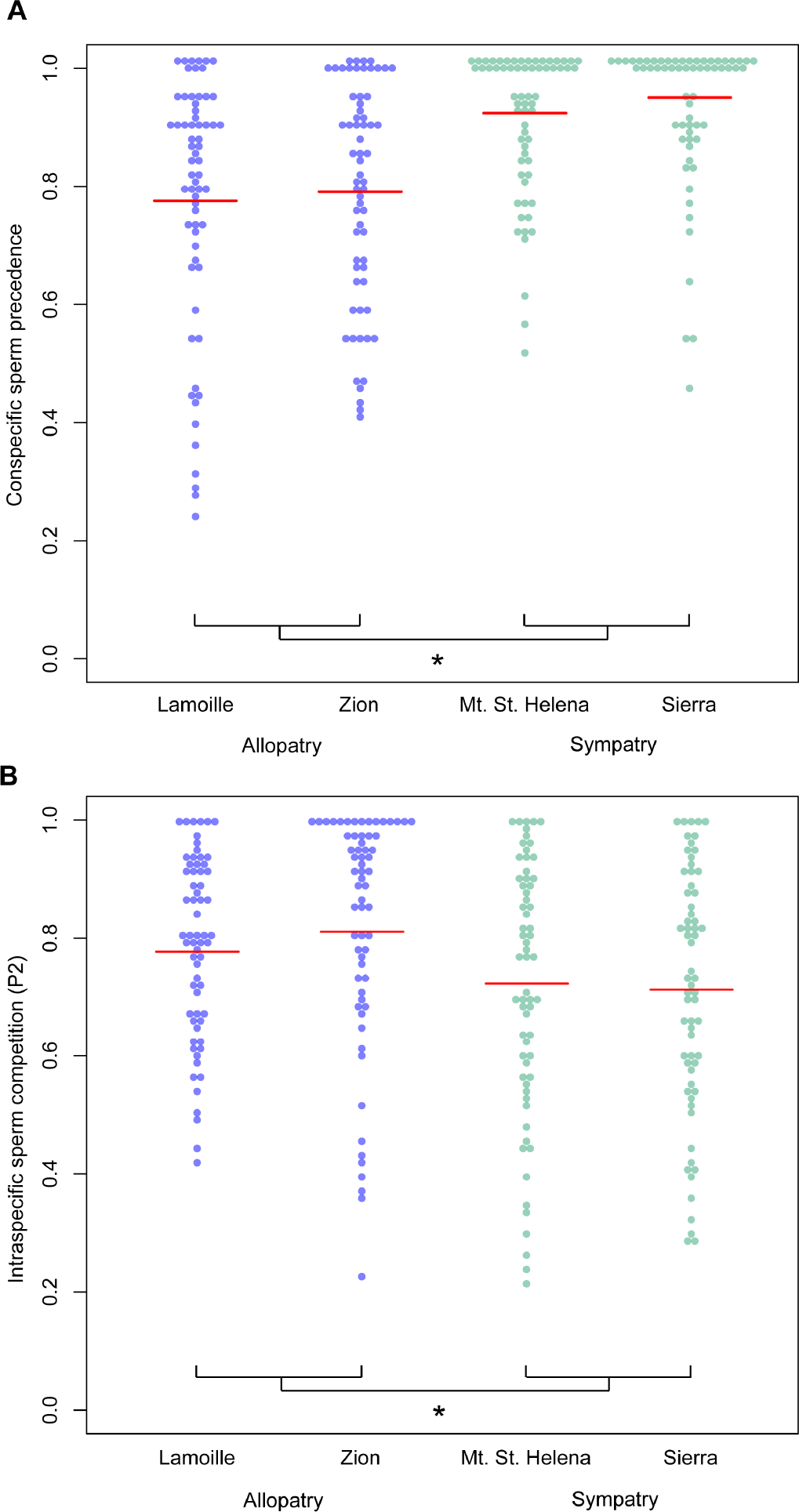
The phenotypic distributions of CSP (panel A) is consistent with a pattern of reinforcement. The distribution of ISC (panel B) shows a shift in ISC in the opposite direction compared to CSP for sympatric populations. The red line in each distribution represents the mean value. Significant differences determined by Welch’s t-test and Wilcox tests between the allopatric and sympatric populations is denoted by *.

### Reinforcement has collateral effects on intrapopulation sperm competition

ISC also differed between allopatric and sympatric populations, in both mean and variance (Table 1; Fig 2B). First, mean offensive ability for ISC was significantly lower in sympatric populations (*t*=3.738, df=246.55,*P*=0.0002; Wilcox’s *W*=10280, *P*=0.0004). This contrasts with the observed increase in offensive CSP in sympatric populations. Second, there was more variation in ISC in the sympatric populations compared to the allopatric populations (Leven-type test χ^2^=5.74, *P*=0.0172). Given the differences in ISC and CSP across populations, we used the mean CSP and ISC phenotype for each male x female genotype combination within a population (i.e., each cell within the diallel crossing design) to examine the pattern of relationship between the two phenotypes across the four populations. We observed a significant negative relationship between CSP and ISC (Pearson’s *r*=−0.31, *P*=0.01; (Fig. 5). Since each male or female genotype is represented in multiple combinations we controlled for non-independence using a linear mixed effect model, and confirmed that the negative slope of the relationship was significant as indicated by a confidence interval that did not overlap zero (Profiled CI = −0.451, −0.028).

### Female genotype effects contribute to CSP and male x female genotype effects explain both CSP and ISC

Of male, female, and male x female genotype effects that could contribute to explaining the variance in CSP, we found that three out of the four populations had a significant female genotype effect on CSP (Table 2; Fig 3), and all populations had a significant male x female genotype interaction effect. The *D. persimilis* tester male line was also significant in three out of four populations. There was no consistent pattern among populations in which effect had the largest intraclass correlation (i.e. which explained the largest proportion of variance; see Methods); in some populations the female genotype effect had the largest intraclass correlation, while in others the male x female genotype interaction had the largest intraclass correlation (Table 2). In contrast, for ISC in all four populations we only observed significant male x female genotype interaction and a significant effect of the first-tester male genotype (Table 3; (Fig. 4). In every case, the male x female genotype effect had a larger intraclass correlation (usually two to three times greater) than the identity of the specific tester male genotype within each *D. pseudoobscura* population.

**Table 2.** The genotype effects that predict CSP. The maximum likelihood estimate (ML est.) and intraclass correlation (ICC) are reported as point estimates from the full model. The *P*-value for each term was calculated by comparing the observed Likelihood ratio test statistic (LR) to the distribution generated by parametric bootstrap. Data were bootstrap sampled according to the null hypothesis where the random effect of interest is not included. The full and reduced models are then fit to each bootstrap sample to determine the distribution for the LR test statistic. A = allopatric; S = sympatric. Bold indicates significance at *P*<0.05. Italics indicates marginal significance *P*<0.06.

|  |  |  |  |  |
| --- | --- | --- | --- | --- |
| Lamoille (A) |  |  |  |  |
| Effect | ML est. | LR | *P*-Value | Intraclass Corr. |
| Female | **0.4024** | **8.10** | **0.0067** | 0.096 |
| Male | 0.0000 | 0.00 | 0.7509 | 0.00 |
| M x F | **0.1154** | **3.52** | **0.0383** | 0.027 |
| *D. persimilis* | **0.3413** | **37.49** | **0.0013** | 0.082 |
| Zion (A) |  |  |  |  |
| Effect | ML est. | LR | *P*-Value | Intraclass Corr. |
| Female | *0.2683* | *2.72* | *0.05632* | 0.067 |
| Male | 0.0000 | 0.00 | 0.4190 | 0.00 |
| M x F | **0.3315** | **16.30** | **0.00238** | 0.0833 |
| *D. persimilis* | **0.0865** | **6.44** | **0.0068** | 0.0217 |
| *Mt St. Helena*(S) |  |  |  |  |
| Effect | ML est. | LR | *P*-Value | Intraclass Corr. |
| Female | **0.8408** | **5.77** | **0.0068** | 0.188 |
| Male | 0.0000 | 0.00 | 0.9891 | 0.000 |
| M x F | **0.3266** | **8.76** | **0.0026** | 0.0737 |
| *D. persimilis* | 0.0000 | 0.00 | 0.9851 | 0.000 |
| *Sierra* (S) |  |  |  |  |
| Effect | ML est. | LR | *P*-Value | Intraclass Corr. |
| Female | 0.3287 | 0.72 | 0.1760 | 0.071 |
| Male | 0.1529 | 0.27 | 0.2673 | 0.033 |
| M x F | **0.5975** | **7.28** | **0.0046** | 0.129 |
| *D. persimilis* | **0.2487** | **8.16** | **0.0012** | 0.053 |

**Figure 3.**
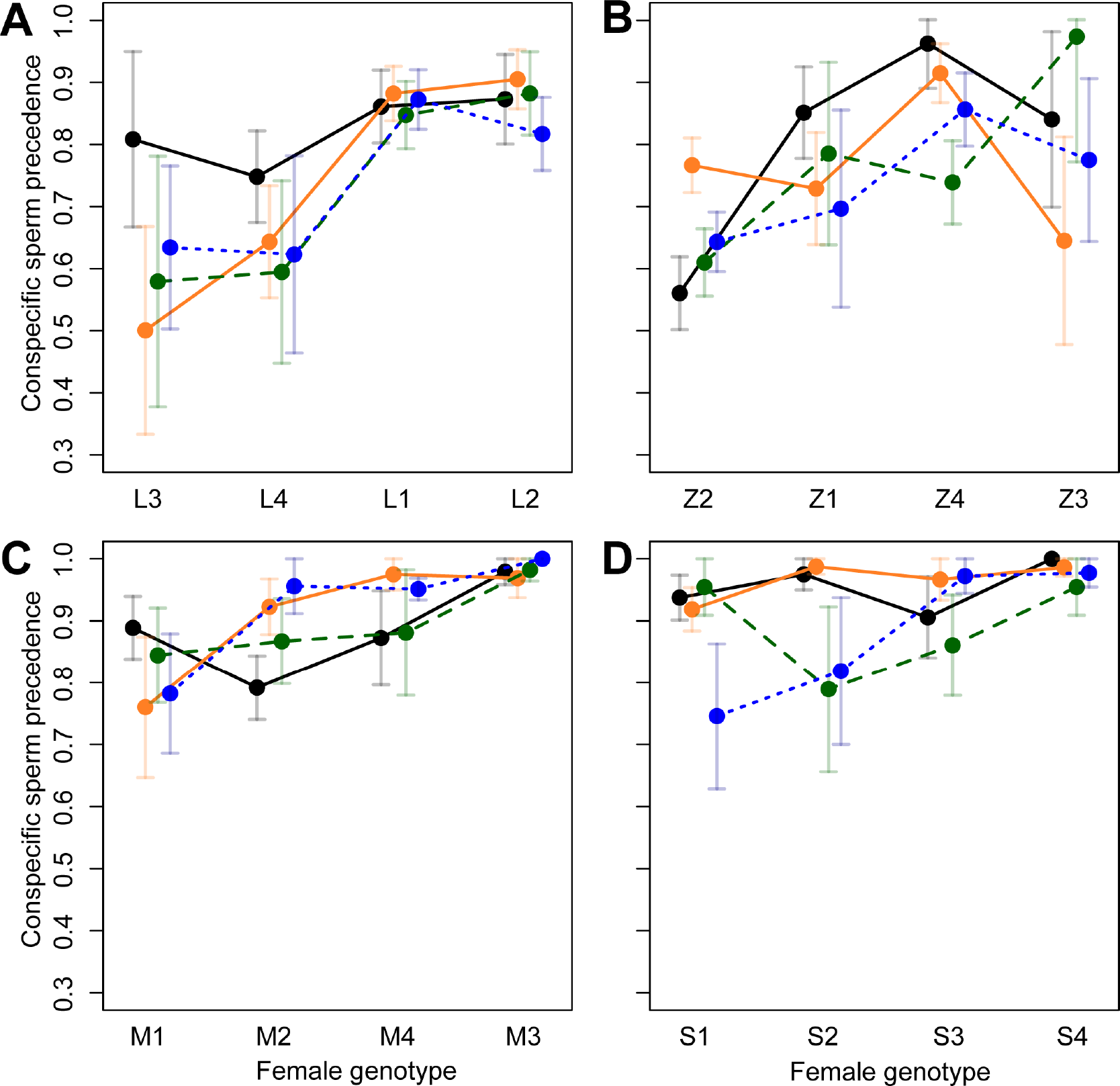
Conspecific sperm precedence (CSP) for all male-female genotype combinations in each population demonstrating a significant effect of female genotype and male-female genotype interaction on the outcome of CSP. A) Lamoille-Allopatry, B) Zion-Allopatry, C) Mt. Dt. Helena-Sympatry, and D) Sierra-Sympatry. Each point represents a specific male-female genotype combination. Error bars are ± one standard error. Female genotypes are ordered by mean CSP. Each color represents a single male genotype for each population. Colors were re-used between each population panel, but actual second male genotypes were unique to each population.

**Table 3.** The genotype effects that predict ISC. The maximum likelihood estimate (ML est.) and intraclass correlation (ICC) are reported as point estimates from the full model. The *P*-value for each term was calculated by comparing the observed Likelihood ratio test statistic (LR) to the distribution generated by parametric bootstrap. Data were bootstrap sampled according to the null hypothesis where the random effect of interest is not included. The full and reduced models are then fit to each bootstrap sample to determine the distribution for the LR test statistic. A = allopatric; S = sympatric. Bold indicates significance at *P*<0.05.

| Lamoille (A) |  |  |  |  |
| --- | --- | --- | --- | --- |
| Effect | ML est. | LR | *P*-Value | Intraclass Corr. |
| Female | 0.0668 | 0.825 | 0.2131 | 0.018 |
| Male | 0.0000 | 0.000 | 0.5037 | 0.000 |
| M x F | **0.2098** | **29.93** | **0.0023** | 0.057 |
| GFP male | **0.0879** | **23.88** | **0.0010** | 0.024 |
| Zion (A) |  |  |  |  |
| Effect | ML est. | LR | *P*-Value | Intraclass Corr. |
| Female | 0.3003 | 3.202 | 0.0647 | 0.074 |
| Male | 0.0405 | 0.170 | 0.3721 | 0.010 |
| M x F | **0.3056** | **22.47** | **0.0022** | 0.076 |
| GFP male | **0.0835** | **12.21** | **0.0011** | 0.020 |
| Mt. St. Helena (S) |  |  |  |  |
| Effect | ML est. | LR | *P*-Value | Intraclass Corr. |
| Female | 0.0000 | 0.000 | 1.0000 | 0.00 |
| Male | 0.0184 | 0.096 | 0.4120 | 0.005 |
| M x F | **0.2195** | **52.44** | **0.0019** | 0.060 |
| GFP male | **0.0825** | **35.24** | **0.0010** | 0.022 |
| Sierra (S) |  |  |  |  |
| Effect | ML est. | LR | *P*-Value | Intraclass Corr. |
| Female | 0.0000 | 0.000 | 0.3744 | 0.00 |
| Male | 0.0000 | 0.000 | 1.0000 | 0.00 |
| M x F | **0.4139** | **70.85** | **0.0021** | 0.111 |
| GFP male | 0.0077 | 0.902 | 0.0886 | 0.002 |

**Figure 4.**
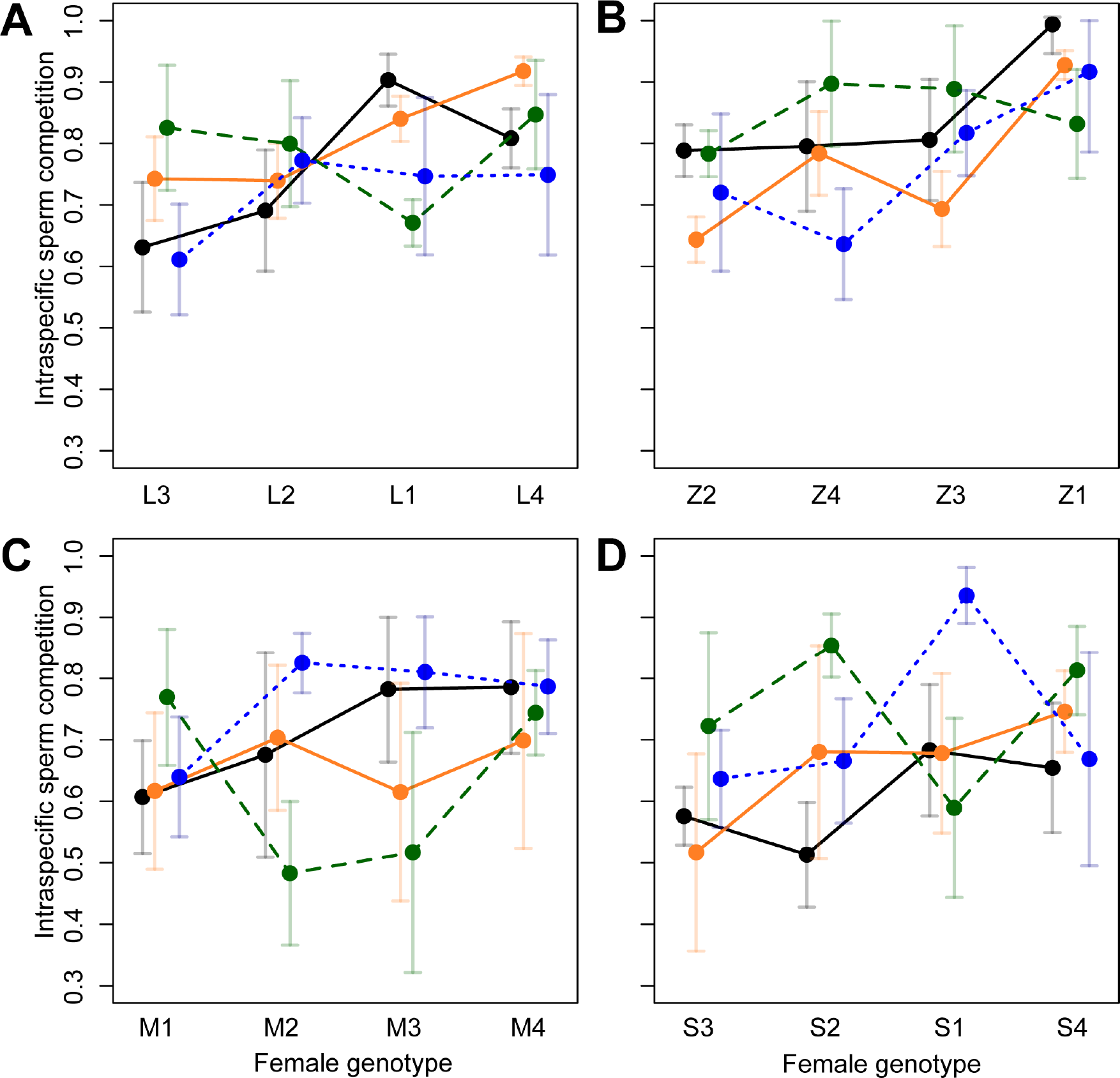
Intrapopulation sperm competition (ISC) for all male-female genotype combination in each population demonstrating a significant male-female genotype interaction on the outcome of ISC. A) Lamoille-Allopatry, B) Zion-Allopatry, C) Mt. Dt. Helena-Sympatry, and D) Sierra-Sympatry. Each point represents a specific male-female genotype combination. Error bars are ± one standard error. Female genotypes are ordered by mean ISC. Each color represents a single male genotype for each population. Colors were re-used between each population panel, but actual second male genotypes were unique to each population.

**Figure 5.**
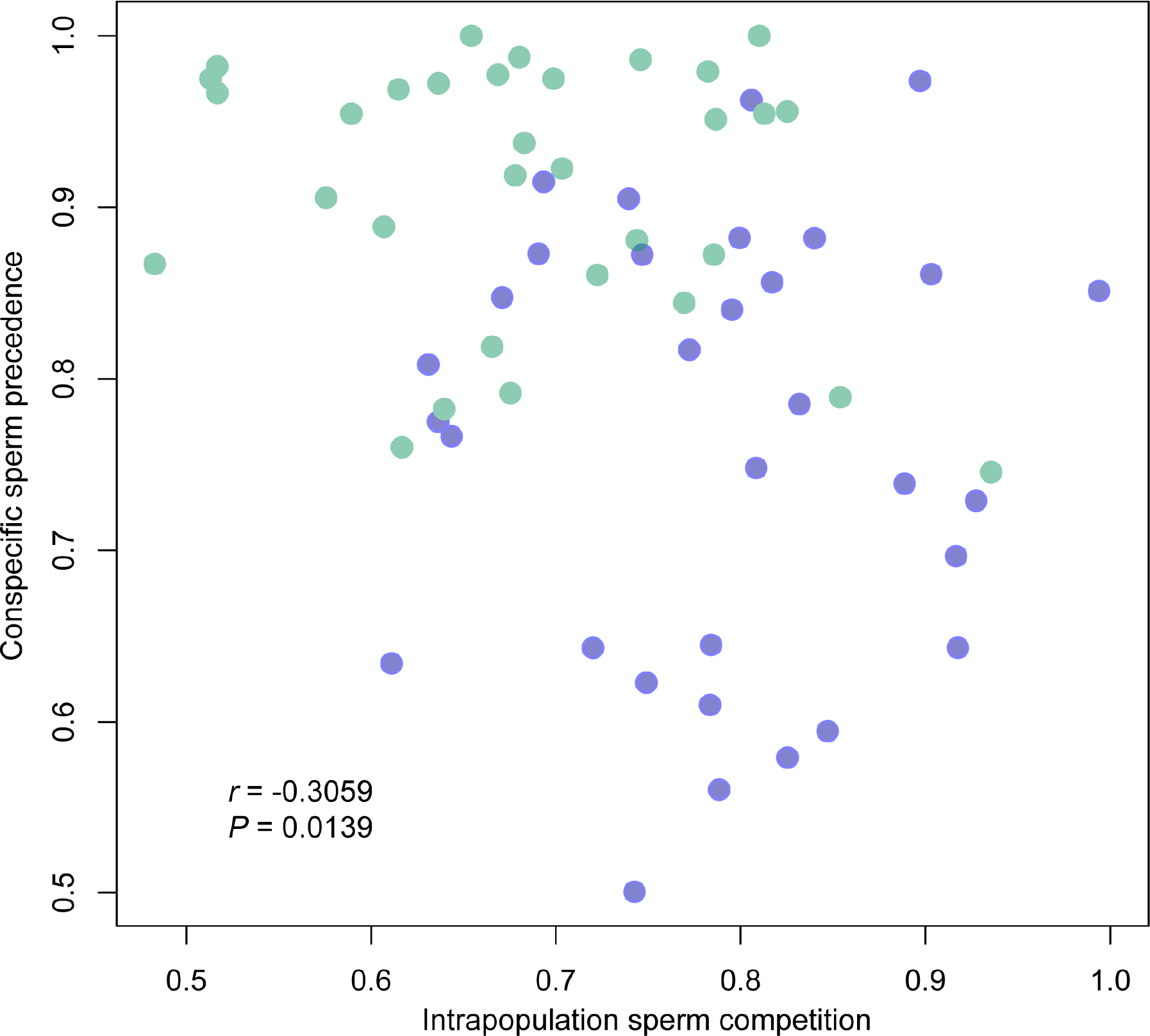
The negative correlation between intrapopulation sperm competition (ISC) and conspecific sperm precedence (CSP) across all four populations with each point representing a male-female genotype combination. Blue points are from allopatric populations and green points are from sympatric populations.

### The opportunity for sexual selection is decreased in sympatry

Our design allowed us to describe the reproductive success of males in terms of offensive (second male) and defensive (first-tester male) success. We found that the sympatric populations had significantly lower variance for reproductive success compared to the allopatric populations (Figure 6; Supplemental Table 4). The variance in reproductive success across all male genotypes (both offensive and defensive) in the allopatric Lamoille population was significantly greater than both sympatric populations (Mt. St Helena *F*=1.96, Bootstrap *P*=0.003; Sierra *F*=2.08, Bootstrap *P*=0.008), as was the variance in reproductive success in the allopatric Zion population compared to the sympatric populations (Mt. St Helena *F*=2.65, Bootstrap *P*=0.003; Sierra *F*=2.83, Bootstrap *P*=0.004).This reduced variance in reproductive success in sympatry is a product of lower offensive sperm competition values in sympatry, that result in equalized differences in the siring success between offensive and defensive males.

**Figure 6.**
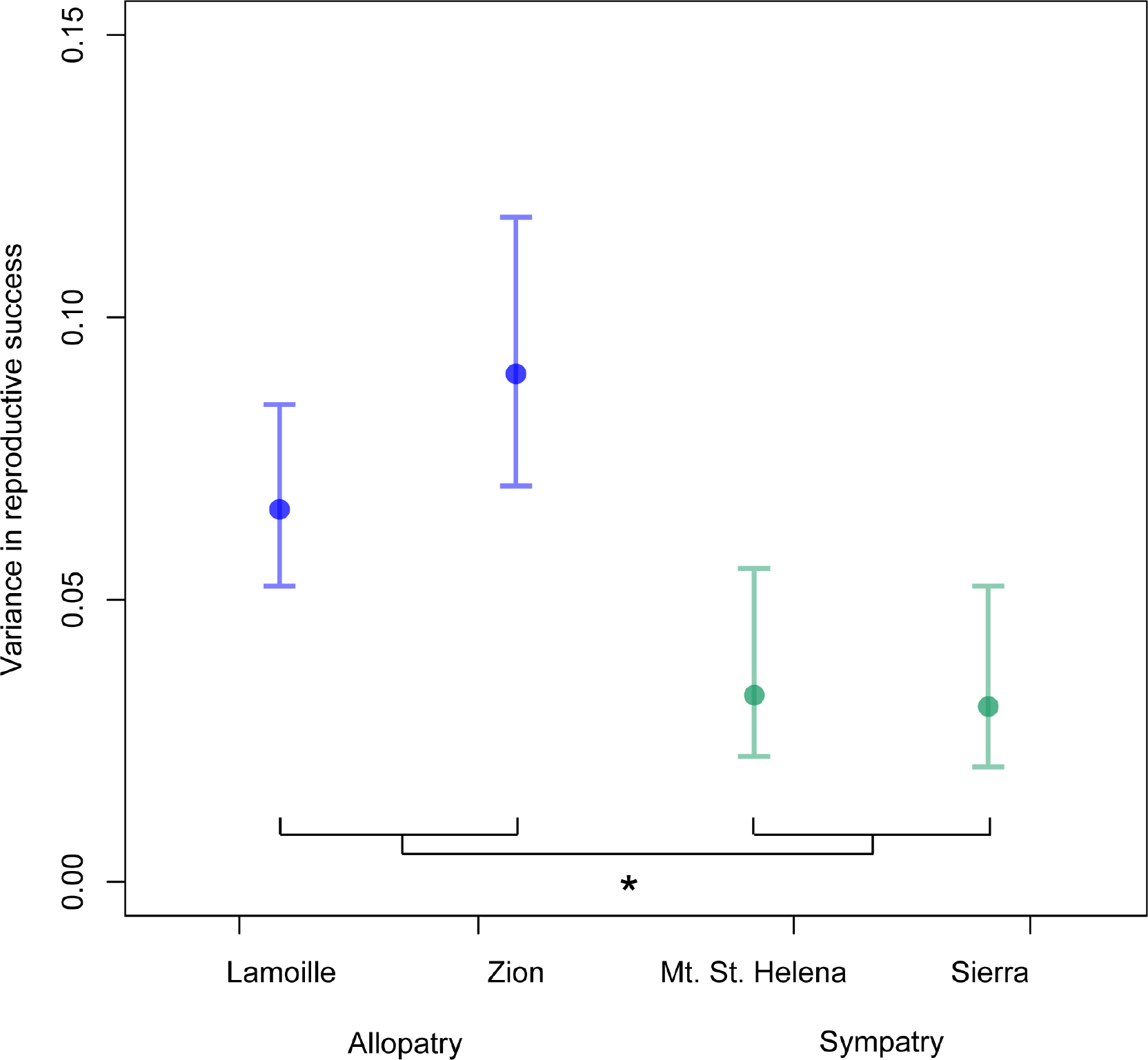
The variance in reproductive success across populations calculated in the framework that combines offensive and defensive males. Each point represents the estimate for the variance in fitness for each population. The error bars are confidence intervals generated from the empirical bootstrap distribution. Significance, denoted by *, was assessed in pairwise comparisons between allopatric and sympatric populations using empirical bootstrap hypothesis testing.

## DISCUSSION

Interactions with heterospecifics have the potential to drive divergent sexual selection and the evolution of reproductive isolation, via reproductive character displacement and reinforcement [6–7,44]. Using *D. pseudoobscura* and *D. persimilis*, we assessed whether there was evidence for reinforcement of species barriers in sympatry via elevated female preference or conspecific sperm precedence, traits that are known to contribute to reproductive isolation across numerous taxa [2]. Premating isolation is historically considered to be a strong barrier to isolation between these species, and one that reinforcing selection has acted on [43], but we saw no evidence for reproductive character displacement for this trait. In contrast we saw a clear signal of increased CSP in sympatric populations, consistent with a pattern of reinforcement. Specifically, the average CSP was higher, and the overall level of phenotypic variation was lower, in sympatric populations, a pattern consistent with recent or recurrent directional selection acting on CSP in these populations. We further asked whether reinforcement could have collateral effects on intraspecific sperm competition and sexual selection, given that these two traits are mechanistically and genetically linked [11,45]. We found that sympatric populations also had lower ISC ability (lower offensive ability) than allopatric populations, consistent with weakened sexual selection in sympatry.

Our results indicate that CSP can strongly contribute to reproductive isolation in response to reinforcing selection. While CSP is known to be a barrier to gene flow in *Drosophila* [12–13] and other taxa [2], its overall importance in nature has been difficult to ascertain [14,16]. Moreover, previous studies of reinforcement sometimes qualitatively describe variation in the target premating traits, but trait variance is typically not quantified [5,9,17] even though models of speciation by sexual selection predict that strong divergent selection will erode phenotypic variation in selected traits [46–47]. Our observations of both increased mean CSP and reduced variation specifically in sympatry provide compelling support for the inference that CSP has responded to strong selection imposed by heterospecific interactions, and underscores the important role that CSP can play in maintaining species boundaries.

The pattern of reproductive character displacement that we observed for CSP is consistent with reinforcement, but other factors have been proposed to account for reproductive character displacement including differential fusion [48] or ecological differences that have collateral effects on mating traits [44,49]. Differential fusion predicts that strong reproductive isolation evolves between species in allopatry and merely prevents species collapse upon secondary contact, so that sympatric species incidentally appear to have stronger isolation [50–51]. If differential fusion operates at the deme/lineage level within a population we would expect the sympatric CSP values to be a subset of allopatric CSP values [17]. This is not the case, however, because the sympatric values of CSP are systematically higher than in allopatry ((Figure 2). Regardless, the differential persistence of demes/lineages with strong CSP in sympatry would nevertheless be consistent with selection from standing variation leading to reinforcement [52,53]. Similarly, several lines of evidence argue that systematic ecological differences between allopatry and sympatry are unlikely to explain our observed postcopulatory differences. Although both sympatric populations are located in California, they are ecological distinct (collected from two different mountain ranges) in terms of numerous ecological factors [54]. Indeed, habitat variation between sympatric populations of *D. pseudoobscura* has led to differences in inversion frequencies maintained by ecological forces that differ in these locations [55]. Moreover, the ecological differences across the whole range of *D. pseudoobscura* are largely continuous, rather than uniquely differentiating regions of allopatry and sympatry/co-occurrence with *D. persimilis*. Given the ecological diversity between populations we do not expect a consistent direction of natural selection acting on either the sympatric or allopatric populations. Arguably more important, there are no established mechanisms whereby external ecological factors are expected to have a direct effect on the strength of sperm competition consistent with our observed pattern. Indirect effects of diet and nutrition can affect sperm competition outcomes [56–57], but should not persist in the lab environment. Moreover, if ecological mechanisms existed there is no reason to expect they would act in the specific direction we observed here. Given this, while the ecological alternative to reinforcement might be plausible for some premating phenotypes, it is unlikely to explain the postcopulatory phenotypes that we examine here.

Our second major inference is that the response to reinforcing selection observed in CSP has had a collateral effect on the magnitude of offensive ISC and the opportunity for sexual selection in sympatric populations. The decrease in the opportunity for sexual selection in sympatry appears to be the result of a negative genetic correlation between CSP and ISC, as well as reduced variance in post-copulatory fitness based on ISC estimates. Sperm competition strongly contributes to sexual selection in *D. pseudoobscura* where multiple mating is frequent in wild caught females [58], and male mating success, including sperm competition, is a major component of selection in natural populations [59]. The observed reduction in offensive sperm competition differs from both of our *a priori* expectations. One *a priori* hypothesis was that selection for increased CSP in sympatry would select for increased offensive sperm competitive ability among conspecifics, if offensive ability were a general trait that acted regardless of whether the competitor was a conspecific or heterospecific male. In contrast, we observed that ISC, was lower for sympatric populations compared to allopatric populations; that is, average offensive ability was closer to 0.5, indicating a greater equalization in sperm competitive ability among competing males. Our other *a priori* expectation was that strong directional selection would alter sexual selection by reducing phenotypic variation. However, the reduced phenotypic variation seen for CSP in sympatry was not mirrored by reduced phenotypic variation for ISC. This observation is also inconsistent with an alternative explanation-that selection for weaker ISC in sympatry indirectly increased CSP. This alternative is more generally implausible as it requires that there has been selection specifically to reduce ISC, solely in sympatry. Instead, we infer that selection for stronger CSP in sympatry has reduced mean ISC in sympatric populations via a negative genetic correlation between these two sperm competitive phenotypes.

For reinforcing selection to influence and interfere with sexual selection, the selection favoring increased CSP must outweigh selection acting to maximize ISC. One way CSP could have a larger effect on fitness than ISC is via a higher selective premium specifically for females. Weaker CSP results in substantial fitness deficits for females because of reproductive investment in low or no fitness hybrids, whereas weaker ISC likely has a comparatively marginal effect on female fitness outcomes. Regardless, the strength of reinforcing selection on CSP depends on the frequency of heterospecific matings. Several lines of evidence suggest that heterospecific mating rates are common between these species. First, from our data we observe a large range in the frequency with which *D. pseudoobscura* females accept *D. persimilis* males in no-choice experiments, with some genotypes on average accepting *D. persimils* 90% of the time. Second, while no estimates for heterospecific mating rate exist from natural populations, rare F1 progeny have been identified from wild collections [60]. Third, genetic evidence suggests there has been post-speciation gene flow (i.e., evidence of movement of alleles between species) between *D. pseudoobscura* and *D. persimils* [61–62]. Notably, these estimates of realized gene flow will systematically underestimate the rate of heterospecific matings, because they will only capture events that result in F1 progeny that themselves then successfully reproduced; for example, given the presence of strong CSP, many heterospecific matings may never produce hybrid progeny.

We were able to test the hypothesis that females face more costs of hybridization [39–41] and that choice manifests as female control of sperm use patterns [63–65] by contrasting the genotype effects (male, female, and male x female genotype effects) between CSP and ISC. We observed significant male x female genotype interactions for all populations for both CSP and ISC but, interestingly, only saw significant female genotype effects for CSP. Significant female genotype effects for CSP suggest that cryptic female choice may be operating similarly to premating isolation mechanisms where females are observed to be the more “choosy” sex and female effects control the level of reproductive isolation more so than male effects [66].

Strong female genotype effects on CSP are also consistent with the current knowledge of postcopulatory sexual selection in the obscura group. Both *D. pseudoobscura* and *D. persimilis* produce two sperm morphs: longer fertilizing eusperm and shorter non-fertilizing parasperm. In *D. pseudoobscura*, the female reproductive tract is spermicidal and higher proportions of parasperm help protect eusperm from these negative effects [67]. Females in sympatric populations may have evolved more effective spermicide against heterospecific males at a cost of spermicidal effectiveness with conspecific males. In this case reproductive isolation would be mediated by cryptic female choice and heterospecific male-female compatibility. This hypothesis may also be consistent with our finding that the *D. persimilis* male genotype contributed significantly to observed variation in CSP.

Reinforcement acting on CSP suggests that other prezygotic barriers that act before CSP are not strong enough to limit the efficacy of selection on CSP in our sympatric populations [14,16]. Indeed, our analysis of premating isolation (propensity to mate with a heterospecific in the first mating) indicated that this potential barrier was equally strong in sympatry and allopatry. This is interesting because one of the first studies demonstrating reinforcement on premating barriers used the *Drosophila pseudoobscura* and *D. persimilis* sister pair [43], although subsequent studies have found more variable patterns [68–70; but see 71]. Our observation of a strong response in CSP also suggests that the populations of *D. pseudoobscura* and *D. persimilis* we examined are not strongly isolated by non-competitive (gametic) isolation, in agreement with inferences from other studies of this specific species pair [70,72]. Though we lack data on CSP from earlier collections in this species pair, our observations here might suggest that the relative contribution of barriers to reproduction has changed in sympatry over time, from premating isolation to CSP. Both gene flow between sympatry and allopatry, or a cost to female premating preferences, might explain this shift over time. Depending on the levels of gene flow among sympatric and allopatric populations, strong premating isolation in sympatry could be lost due to “swamping effects” of allopatric gene flow [73] or could lead to greater species wide reproductive isolation [74–75]. Our data suggest that it’s unlikely that gene flow from sympatry into allopatry created greater reproductive isolation in allopatry (thereby reducing the signal of reinforcement) because the average allopatric premating isolation in our experiment is similar to previous reports [43]. This suggests that reduced premating isolation has emerged in sympatry, but it is difficult to disentangle the effects of gene flow from the cost of female choice as causes of this reduction. Both processes could contribute to the large variance we see for female preference in sympatry compared to the more uniform level of premating isolation in allopatry (Fig 1). The probability that strong female preference have been lost in sympatry also depends on the frequency of this trait and any associated costs of choosiness. When *D. pseudoobscura* stocks are kept in the absence of heterospecific interactions female preference against heterospecifics decreases with longer periods of experimental allopatry, suggesting that it may be costly to maintain this trait [76]. In either case, the reduction in the strength of premating isolation in sympatry suggests that this barrier to reproduction may only generate transient patterns of reinforcement.

Overall, our data suggest that strong reinforcing selection for reproductive isolation can have consequences for sexual selection and sexual interactions, in these important postmating sperm competition traits. The direction of this interaction provides an interesting inversion to standard expectations about the connection between sexual selection and speciation. Sexual selection is often thought of as a driver of sexual characteristics whose evolutionary divergence then contributes to reproductive isolation. But a direct genetic connection between these processes implies reproductive isolation also has the reciprocal potential to shape sexual selection [77]. Based on our observations of higher mean but lower variance in CSP in sympatry, a negative correlation between CSP and ISC, and reduced variance in reproductive success via ISC among sympatric conspecific males, we infer that strong selection for reproductive isolation within populations exposed to heterospecific species has reduced the efficacy of sexual selection in these populations, a collateral effect of reinforcing selection that has not previously been demonstrated.

## ACKNOWLEDGEMENTS

We would like to thank E. Walburn and J. Roesener for their assistance with crosses and scoring progeny, J. Powers and the IU Light Microscopy Imaging Center for assistance with the Leica microscope, M. Noor, A. Hish, and N. Phadnis for providing strains used in this experiment, and Donn Castillo for help with collecting strains. Collections were completed with assistance from IU Biology Department travel awards to DMC. Research was supported by Indiana University Dept. of Biology funding to LCM and an AmericanSociety of Naturalists student research award to DMC. DMC was supported by a President’s Diversity Initiative Dissertation Fellowship from the Indiana University Graduate School.

## MATERIALS AND METHODS

### Wild type fly stocks

All stocks were reared on standard media prepared by the Bloomington Drosophila Stock Center, and were kept at room temperature (~22C). We used a set of isofemale lines collected from four natural populations in the summers of 2013 and 2014. Allopatric D. pseudoobscura were collected at Zion National Park, UT (kindly provided by N. Phadnis) and Lamoille Canyon, NV (collected by D. Castillo). Sympatric D. pseudoobscura and D. persimilis were collected at two sites: Mt. St. Helena, CA (D. pseudoobscura collected by A. Hish/M. Noor and D. Castillo, and D. persimilis collected by D. Castillo); and, near Meadow Vista and Forest Hill, CA (called here ‘Sierra’; D. pseudoobscura and D. persimilis collected by D. Castillo). For both sympatric populations, both species were present in field collections and can be considered truly co-occurring/sympatric.

### Conspecific sperm competition assay

Sperm competition assays generally involve mating an individual female sequentially with two distinct male genotypes. In all experimental crosses between species, females were paired first with a D. persimilis male and second with a D. pseudoobscura male; that is, the assays are evaluating the “offensive” sperm competitive ability of conspecific males to displace heterospecific sperm (equivalent to ‘P2’, or second male siring ability; [78]). We focused on “offensive” sperm competition because D. pseudoobscura females do not remate with D. persimilis males if they have first mated with a conspecific, therefore we cannot evaluate “defensive” sperm competition in this cross. In this experiment we partitioned the variance in CSP due to male genotype, female genotype, and the male x female genotype interaction using a “diallel-like” crossing design, which is commonly used for this purpose [31,33; Supplemental Fig 1). A diallel cross is a mating scheme commonly used to estimate the genetic effects, additive genetic variance, and heritability, of quantitative traits by crossing all parental genotypes in all possible combinations [79]. Our design is “diallel-like” because we did not use progeny from the diallel to estimate heritability. We completed separate CSP experiments for each of our four D. pseudoobscura collection locations (Sympatric= Sierra and Mt. St. Helena, Allopatric= Zion and Lamoille). For each population we used a 4x4x4 design: four D. pseudoobscura female genotypes from that population, four D. persimilis genotypes as first males (“tester males”), and four D. pseudoobscura male genotypes as second males from the same population as females. Each 4x4x4 combination was replicated once (n=64 unique cross combinations for each population). If CSP is important for reproductive isolation in sympatry it should be consistently strong across multiple heterospecific genotypes. Accordingly, rather than rely on a single D. persimilis genotype, we aimed to use multiple wild-collected D. persimilis tester male lines for our experiments. Of these, two D. persimilis lines were collected at the same time and in the same traps as the D. pseudoobscura strains at the Sierra location and another two at the Mt. St. Helena location.

Virgin individuals were collected and aged 7 days prior to the initiation of an experimental block. One day before mating, D. persimilis tester males were isolated individually [80]. The following day, females were individually added (without anesthesia) to a vial containing a tester male and were co-housed for 24 hours, after which time the tester male was removed. We kept females housed individually in these vials for 7 days before second mating (similar to [80]). After 7 days we inspected all vials for the presence of larvae to determine if females had mated with the first D. persimilis tester males. This was used to evaluate evidence for differences in successful first matings (pre-mating isolation) among allopatric and sympatric populations, rather than observing matings directly, as there is high variance in time to copulation in this heterospecific pairing [70]. Only females that had mated (i.e. had produced larvae within 7 days) were retained for the remainder of the CSP experiment.

For the second mating, each individual female was paired with one of the four D. pseudoobscura male genotypes from her own population to determine the strength of CSP. These second males were also isolated one day before the introduction of the female. Seven days after mating with the first male, females were transferred, without anesthesia, to the vial containing the second male. Individual pairs were co-housed for 24 hours and the male was removed on the second morning. The female was kept for five days (transferring after 2 days to avoid overcrowding of larvae). All progeny produced in the five-day window after the second mating were collected; from these progeny a maximum of 10 males and 10 females, randomly chosen from the total group of progeny, were used to score CSP (P2) as described below.

### Intrapopulation sperm competition assay

The design for intrapopulation sperm competition (ISC) assay mirrored the experimental design for CSP except that, rather than a D. persimilis tester male, the first male was a D. pseudoobscura tester male derived from the same population as the D. pseudobscura female and second male genotypes in the trial. For each population we used a 4x2x4 design: four D. pseudoobscura female genotypes, two D. pseudoobscura GFP genotypes as first males and 4 D. pseudoobscura male genotypes as second males. The same female x second male genotypes were used in ISC and CSP experiments. Each combination was replicated twice (n=64 for each population, with 32 unique cross combinations). This allowed us to have a total sample size per population that matched the CSP experiment (64 replicates per population, 256 replicates across all populations).

The details of the mating scheme (virgin collection, aging of individuals, isolation of individuals, etc.) are identical to the CSP experiment. We did not observe matings directly, but the average refractory period for D. pseudoobscura is 4 days [81], so we are confident that on average only a single mating occurred in the 24 hour co-housing timeframe. Each individual female was randomly assigned one of the two D. pseudoobscura first male (tester) genotypes to determine the strength of P2 (second male siring ability) by our four focal second male genotypes, against these tester male genotypes. The female was kept for five days after the second mating (transferring after 2 days to avoid overcrowding of larvae). All progeny produced in the five-day window after the second mating were collected and scored.

### Generating visibly-marked tester males for quantifying CSP and ISC

To allow efficient progeny scoring, paternity was scored with the aid of visible markers in both CSP and ISC experiments. This required us to generate marked male tester lines with wild-caught D. persimilis (for CSP tester males) and D. pseudoobscura (for ISC tester males) lines from each study population.

For CSP, to introduce a visible marker into wild-type wild-collected D. persimilis males from our sympatric sites, we introgressed an X-linked marker (“short” or sh) from a D. pseudoobscura line, into four of our collected D. persimilis genotypes (Supplemental Fig 2). These four D. persimilis tester males originated from isofmale lines collected at the Sierra and Mt St. Helena locations and were used to evaluate the mean strength and variation in CSP for all four D. pseudoobscura populations in the CSP experiment. We first crossed these D. persimilis sh mutant males to females from each of the four wild-type D. persimilis isofemale lines (keeping each tester genotype separate throughout this process). This produced F1 daughters heterozygous for the sh allele, that were backcrossed to wild type males from the same wildtype isofemale line. From the BC1 progeny we retained sh males, and these were backcrossed to the original D. persimilis isofemale line to generate BC2s (Supplemental Fig 2). This process of alternating males and females for each backcross generation within each D. persimilis isofemale line was completed until the BC12. The alternation of male/female during backcrossing was necessary because recombination only occurs in females, but to retain the marker we had to select for sh males every second generation. After the BC12, the progeny within each BC isofemale line were interbred to create males and females homozygous for the sh allele. We did not directly evaluate how much of the sh line genome was introgressed in each case, however, D. pseudoobscura and relatives have a much higher recombination rate than D. melanogaster [82], and previous introgression lines between these species have eliminated unwanted regions after 4 generations of backcrossing [83].

For ISC experiments, the marked tester males were created by introgressing a green fluorescent protein marker (GFP) into 2 wild type D. pseudoobscura strains per location (therefore 8 strains in total, using wild-collected isofemale lines that were not used as female or second male genotypes for the ISC experiments). The original GFP strain was obtained from the UCSD stock center (14011-0121.166) the creation of which is described in Holtzman et al. [84]. We chose this marker because it is dominant [11]. We chromosomally mapped the GFP insertion of the original GFP strain to the second chromosome using a multiply marked (MM) strain, which contains visible recessive markers on all of the major chromosomes (y;gl;or;inc kindly provided by N. Phadnis, University of Utah). This mapping was completed in order to ensure the GFP insertion was not on the 3rd chromosome which, in D. pseudoobscura, contains large inversions that would have inhibited recombination of the marker into the wild-type backgrounds of our D. pseudoobscura isofemale lines.

The original GFP line was created in a stock that carried the X-linked white mutation. To eliminate the white allele from the population, in the parental cross we crossed the WT line with the GFP carrying male, and then used only F1 males with wild-type X chromosomes (no white mutation) to backcross in this initial generation. For the remaining eight backcross generations, we used females to allow recombination. We then chose 10 sibling pairs for each genotype to ensure the GFP marker was homozygous. These sub-lines were inbred for two generations. In the second generation we testcrossed the founder pair of individuals of each sub-line to ensure they were homozygous for the GFP marker. We recovered 2-4 lines that were homozygous for the GFP marker for each genotype. We then combined inbred lines that had originated from the same isofemale genotype to reduce any potential effects of inbreeding depression that might have arisen during marker introgression.

### Scoring conspecific sperm precedence

Hybrid male progeny from D. pseudoobscura x D. persimilis crosses are sterile (there are no motile sperm, observable by dissecting the testes). We used this sterility phenotype to differentiate the male progeny of heterospecific versus conspecific males and therefore to score CSP. For a given replicate we collected and dissected 10 male progeny that were produced after the second mating. Each male was dissected individually in PBS buffer, and its testes moved to a slide that had 1ul of PBS buffer. A cover slip was placed over the slide and the testes were squashed, releasing sperm into the buffer. The slides were examined under an EVOS FL microscope for the presence of motile sperm. If no motile sperm were present, the male was scored as hybrid.

Because female hybrids are fertile in these crosses, the sh allele was used to differentiate the female progeny of heterospecific versus conspecific males and therefore to score CSP from female offspring. Since the sh allele is recessive we could not score F1 females directly, but instead scored their offspring for the presence of the sh allele. If an F1 female was hybrid (and carrying the sh allele from the D. persimilis male) we would expect half of her sons and half of her daughters to have the sh phenotype. We previously confirmed that the half segregation held for known hybrid progeny. For each cross, ten F1 females (that could be hybrid or purebred) were housed individually with a D. pseudoobscura male that also carried the sh allele (UCSD stock center Dpse co;sh 14011-0121.13). We chose a D. pseudoobscura male for these crosses to increase the number of progeny to score since D. pseudoobscura females (and therefore any purebred female progeny in our experiment) exhibit premating isolation with D. persimilis males; hybrid females do not demonstrate a mating preference. After a week the parental individuals were cleared from the vials and the vials were retained to score progeny. As progeny eclosed they were scored for the presence of sh allele. Any F1 female that produced sh progeny was considered hybrid. We required each F1 female to produce at least 10 progeny to be used in scoring CSP.

Our measure of CSP was then the number of purebred progeny out of the total number of F1 individuals scored for a particular cross. If all progeny produced in a cross were scored as hybrid, we did not use this replicate in our analyses because we could not ensure that a second mating had taken place. Note that the frequency of this failure to remate following a first mating does not differ between populations [70]. Every CSP estimate was based on at least 10 scored progeny and, for the majority of the crosses, we scored close to 20 individuals. In addition, to ensure that CSP estimated here does not simply reflect stronger fecundity stimulation by conspecific males, in a pilot experiment we determined that there was no difference in progeny production in heterospecific vs. conspecific matings for any of the allopatric or sympatric populations, consistent with previous work [70,72]. There was also no correlation between the total number of progeny scored for CSP and the magnitude of CSP, and the number of progeny scored did not differ between populations. These observations suggest that there are no postzygotic survivorship barriers in hybrids between these species that would systematically differ between sympatric and allopatric populations, confounding our estimate of CSP.

### Scoring intrapopulation sperm competition

We scored all progeny that eclosed in the five days after the second mating for the presence/absence of the GFP phenotype. Our measure of sperm competition (P2) for ISC was then the number of wild-type (non-GFP) progeny out of the total number of progeny scored for a particular cross. If all progeny produced in a cross were GFP, we did not use this replicate because we could not ensure that a second mating had taken place. (As with CSP, the proportion of females that did not remate was not significantly different between populations). Individuals were scored as they eclosed, using a Leica M205FA Stereo Microscope that has an Hg fluorescent lamp attached and GFP filter. Individuals were anesthetized and the ocelli were examined for GFP signal as described in Castillo and Moyle [11].

### Statistical analyses

All analyses were completed in R v 3.01.

Differences in the probability of first mating with heterospecifics

We evaluated evidence for a pattern consistent with reinforcement acting on first mating (simple prezygotic isolation) in two ways. First, we used a χ^2^ test of independence to test the null hypothesis that the mating rate with heterospecifics was the same for alternative geographic scenarios (allopatric vs. sympatric), after combining both allopatric and both sympatric populations for this single comparison (pairwise tests among individual populations gave the same result). Second, because χ^2^ tests might lack power, and since mating events can be coded as a binary variable (0 for did not mate, 1 for successful mating), we used a logistic regression model with all four populations represented by a categorical variable using the glmer function. We then tested whether there were any differences in heterospecific mating between populations by conducting a Wald’s test (using the wald.test function from the aod package; [85]).

To evaluate whether there was significant variation within each population (i.e., among isofemale line genotypes) in the probability of mating with a heterospecific, we used logistic regression. We first fit a full model where the probability of mating with a heterospecific depended on the isofemale line, the D. persimilis tester line, and the male x female genotype interaction, and tested significance of these effects using a Wald’s test. Because there was no significant interaction for any population, we fit a reduced model that only contained the effects of isofemale line and D. persimilis tester line without the interaction, and report these models in the results.

### Differences in mean and variance of CSP and ISC between populations

We evaluated evidence for a pattern in CSP consistent with reinforcement, by evaluating whether the allopatric and sympatric populations had a mean difference in CSP or whether they differed in variance. For analyses of mean differences, we pooled the two allopatric populations because there was no significant difference in mean CSP between them (Allopatry *t* = −0.45064, df = 123.62, *P* = 0.653) and pooled the two sympatric populations for the same reason (Sympatry *t* = −0.86678, df = 125.87, *P* = 0.3877). We tested the hypothesis that the mean CSP differed between geographic scenarios using a Welch’s *t*-test that accounts for unequal variances between samples, and (given that the data are not normally distributed) we also confirmed these results with a Wilcoxon ranked sum test. To evaluate differences in variance, we again pooled the allopatric and sympatric populations because the variance was equivalent between allopatric populations (χ^2^ = 0.031899, *P* = 0.8585), and between sympatric populations (χ^2^= 0.80562, *P* = 0.3711). We compared the total phenotypic variation between geographical classes of population with a Levene-type test implemented in the lawstat package in R [86]. The specific test we used in the lawstat is a Kruskal-Wallis modified Brown-Forsythe Levene-type test. The Brown-Forsythe test is based on the absolute deviations from the median, which retains statistical power for many types of non-normal data [87]. Kruskal-Wallis tests are rank-based tests. We used the Kruskal-Wallis modification because the variance in proportion data derived from binomial data does not accurately reflect variance in the original data [88].

Using the same statistical approach as for CSP, we tested for differences in the mean and variance between sympatric and allopatric populations for ISC, again pooling the individual allopatric and sympatric populations as they were not significantly different from one another for either measure (Allopatric mean *t*=−1.136, df=118,66, *P*=0.2593; Sympatric mean *t*=0.191, df=125.72, *P*=0.8488; Allopatric variance χ^2^==0.949, *P*=0.3316; Sympatric variance χ^2^=0.0796, *P*=0.7782). Note that, although we report results from tests with these pooled data in the main text, we also observed significant differences in pairwise tests between individual allopatric and sympatric populations, for both average and variance measures of CSP and ISC (Supplemental Tables 2 and 3).

### Genetic variation and genotype effects on CSP and ISC

Within each population we assessed whether female, male, or female x male genotype predicted variation in the strength of CSP and ISC. While this can be tested using a two-way ANOVA with interaction, the assumptions of ANOVA, including normally distributed residuals and heterogeneity in the distribution of the residuals, are typically violated by binomial data such as our sperm competition data [89–90]. We instead chose binomial regression, as this more naturally models our count/binomial data. The model is of the form

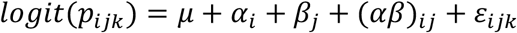

The variable α is a categorical variable with four levels that represents male genotype. The variable β is also a categorical variable with four levels that represents female genotype. The variable (αβ) represents the male x female genotype interactions. Since we were interested in partitioning the variance and estimating the variance components 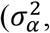 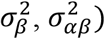 we assumed that each variable was a random variable. To test the significance of each variance component, we used a binomial regression in a mixed modeling framework with parametric bootstrap [91]. In this bootstrap procedure, data are simulated from the null model which lacks the random effect of interest. Then the full and reduced models are fit to the simulated data to determine the bootstrap distribution of the Likelihood Ratio test statistic. To the model above we also included a random effect of tester male (*D. persimilis* for CSP and GFP *D. pseudoobscura* strain for ISC). To provide an assessment of the relative importance of each variable we calculated the intraclass correlation for each coefficient; a high correlation indicates that the variable explains much of the variance in the data. The ICC for the female effect, for example, would be:

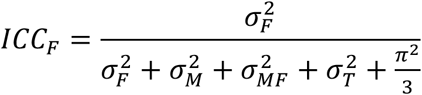

Where *F* represents female variance, *M* represents male variance, *MF* represents the interaction, and *T* represents the identity of the tester male. The 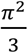 replaces the residual variance for the binomial model with logit link function. In the case of binomial regression the ICC values are on the log scale, and there is no convenient transformation to proportion scale [92], so they are presented here as a relative measure of variance explained.

### Quantifying sexual selection and variance in male reproductive success

To evaluate whether the intensity/opportunity for sexual selection differs among populations we require an estimate of variance in male reproductive success [93]. In a natural population most males can gain fitness through offensive (P1) and defensive (P2) sperm competition, so the best estimate for variance in reproductive success would be total progeny produced. In our experiment we did not score lifetime progeny production, and specific male genotypes were either used as offensive or defensive males only. As such we estimated male fitness as the proportion of progeny sired, taking into consideration that we had two distinct classes of males—tester first (defensive) males and second (offensive) males--that may differ in their frequency and variance in fitness in the experiment. Following Shuster et al. [94] we define total variance in male reproductive success as the sum of within and between male class variance

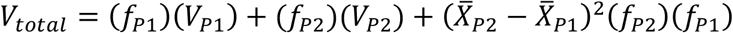

The two terms on the left hand of the equation represent the within class variance (for example, *V*_*P1*_ is the variance in sperm competitive success between tester males and *f*_*P1*_ is the frequency of tester males used in the experiment). The last term represents the between class variance.

We were interested in reproductive variance at the level of male genotype so we averaged biological replicates to generate mean fitness values for each individual genotype. We used empirical bootstrap confidence intervals to estimate error that may have been a product of averaging over replicates [95–96]. For the bootstrap procedure we sampled 16 data points, with replacement, from the 16 original empirical replicates for each genotype (32 for defensive males). We then averaged these data points and calculated *V*_*total*_ as described above. We completed 1000 bootstrap replicates for each population. We constructed the 95% confidence interval using the bootstrap difference 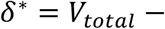 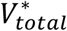 where * represents each bootstrap replicate. The interval is then [*V*_*total*_ − *δ*\*_0.05_, *V*_*total*_ − *δ*\*_0.95_].

The confidence intervals for the Zion population did not overlap with the confidence intervals for either sympatric population and can be considered significantly different at the 0.05 level (Supplemental Table 4). The Lamoille population confidence intervals overlapped with the sympatric populations, but overlap in confidence intervals does not mean parameters are not statistically different [97]. This is because confidence intervals calculated for independent parameters cannot replace a comparative test of the differences between two parameters. Therefore, we conducted bootstrap hypothesis testing [95–96] to determine whether differences in *V*_*total*_ between populations were significant, specifically by calculating bootstrap F statistics. The *F* statistic is a ratio of any two variance parameters, for example 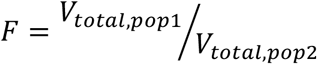. We compared the *V*_*total*_ in pairwise comparisons following standard bootstrap methods, where bootstrap samples are generated under the null hypothesis, and then this distribution is compared to the empirically observed statistic. For our scenario, the null hypothesis was that there was no differences in *V*_*total*_ between populations. Therefore we sampled, with replacement, offensive and defensive genotypes after pooling data from both populations. We then randomly assigned each value to one of the two populations. This generated a bootstrap replicate with approximately equal variance between the populations. We then could calculate 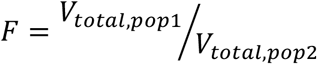 for each replicate. The bootstrap p-value is then calculated by comparing the bootstrap statistic (*F*\*) to the observed statistics (*F*) using 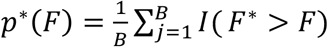. I() is an indicator function that is equal to 1 when the argument is true (boostrap statistic > observed statistic), and 0 when false. B is the number of bootstrap replicates (1000 per population comparison).

